# Restoring adiponectin via rosiglitazone ameliorates tissue wasting in mice with lung cancer

**DOI:** 10.1101/2023.07.31.551241

**Authors:** Henning Tim Langer, Shakti Ramsamooj, Ezequiel Dantas, Anirudh Murthy, Mujmmail Ahmed, Seo-Kyoung Hwang, Rahul Grover, Rita Pozovskiy, Roger J. Liang, Andre Lima Queiroz, Justin C Brown, Eileen P. White, Tobias Janowitz, Marcus D. Goncalves

## Abstract

The cancer associated cachexia syndrome (CACS) is a systemic metabolic disorder resulting in loss of body weight due to skeletal muscle and adipose tissues atrophy. CACS is particularly prominent in lung cancer patients, where it contributes to poor quality of life and excess mortality. Using the Kras/Lkb1 (KL) mouse model, we found that CACS is associated with white adipose tissue (WAT) dysfunction that directly affects skeletal muscle homeostasis. WAT transcriptomes showed evidence of reduced adipogenesis, and, in agreement, we found low levels of circulating adiponectin. To preserve adipogenesis and restore adiponectin levels, we treated mice with the PPAR-γ agonist, rosiglitazone. Rosiglitazone treatment increased serum adiponectin levels, delayed weight loss, and preserved skeletal muscle and adipose tissue mass, as compared to vehicle-treated mice. The preservation of muscle mass with rosiglitazone was associated with increases in AMPK and AKT activity. Similarly, activation of the adiponectin receptors in muscle cells increased AMPK activity, anabolic signaling, and protein synthesis. Our data suggest that PPAR-γ agonists may be a useful adjuvant therapy to preserve tissue mass in lung cancer.

**Key points:**

- The PPAR-γ agonist, rosiglitazone, restores circulating adiponectin levels in mice with lung cancer.
- Rosiglitazone preserves skeletal muscle and adipose tissue mass in mice with lung cancer.
- The preservation of muscle mass with rosiglitazone is associated with increases in AMPK and AKT activity.
- Stimulation of adiponectin signaling increases AMPK activity, anabolic signaling, and protein synthesis in muscle cell culture.

## Introduction

The cancer associated cachexia syndrome (CACS) is a frequently occurring adverse effect of lung cancer, and independently predicts poor quality of life, reduced anti-cancer treatment tolerance, and reduced life expectancy in patients [1, 2]. CACS is characterized by loss of adipose and muscle tissue. It affects up to 50% of patients with lung cancer at diagnosis and up to 80% of patients with advanced disease [3]. There are no clinically established treatment options to prevent tissue wasting and treat CACS. To address this unmet need, we previously characterized a genetically engineered mouse model of lung cancer that closely models human cachexia [4]. LSL-Kras^G12D/+^;Lkb1^f/f^ (KL) mice develop lung tumors that result in the CACS phenotype in 80% of the mice, while the remaining mice maintain their body weight, muscle mass, and fat mass [4].

The CACS phenotype in humans with lung cancer and KL mice is characterized by severe atrophy of the white adipose tissue (WAT) [5]. WAT is not only the body’s reservoir for energy-dense metabolites and lipids, but it also serves as a key energy sensor, actively communicating with other organs and tissues to maintain energy balance. This communication is accomplished by the production and secretion of a variety of hormones, including leptin and adiponectin. These hormones help to regulate body weight by controlling food intake, energy expenditure, and insulin sensitivity. For example, adiponectin improves insulin sensitivity and enhances glucose uptake and utilization in skeletal muscle, which contributes to the maintenance of metabolic homeostasis in the body.

We have previously utilized the incomplete penetrance of CACS in KL mice to identify targetable pathways in the liver and tumors by performing comparative transcriptomics on tissues from mice with and without CACS [4, 6]. In this study, we used the same approach on WAT samples to find evidence of reduced adipogenesis and adiponectin secretion in mice with CACS. We targeted adipogenesis and restored levels of circulating adiponectin in KL mice with the PPAR-γ agonist, rosiglitazone. Rosiglitazone treatment delayed weight loss and preserved WAT and skeletal muscle mass without changes in tumor burden. These beneficial effects were associated with enhanced AMPK signaling in skeletal muscle. Furthermore, we found that stimulation of the adiponectin receptor results in dose-dependent activation of AMPK, improved anabolic signaling, and enhanced muscle protein synthesis in muscle cell culture. These data suggest that WAT dysfunction contributes to the loss of skeletal muscle mass during CACS, and PPAR-γ agonists may be a useful adjuvant therapy to preserve tissue mass in subjects with lung cancer.

## Results

### Cancer cachexia features impaired adipogenesis and adiponectin expression in mice

Adipose tissue and skeletal muscle wasting are the most common signs of CACS. However, the interaction between these tissues is poorly understood. To better comprehend how loss of adipose tissue impacts muscle physiology during CACS, we performed RNA-Seq on WAT tissue obtained from KL mice with and without CACS [4]. Distinct transcriptomic changes were obvious from a principal component analysis (PCA), which revealed differential clustering of the mice with and without CACS (Figure 1A). We next performed a Gene Set Enrichment Analysis (GSEA) using the Hallmark database to identify differentially expressed pathways. The top upregulated pathways included the estrogen response, KRAS signaling, and the P53 pathway (Figure 1B). The top downregulated pathways included E2F targets, G2M checkpoint, and adipogenesis. We previously found that KL mice with CACS have less WAT mass, smaller adipocytes, and lower production of leptin [6]. While leptin is not a direct marker of adipogenesis, its circulating levels can be used as an indicator of adiposity and potentially as a biomarker for adipocyte size [7]. We assessed the levels of another major adipocyte-derived hormone, adiponectin, because circulating levels typically rise when adipose tissue is depleted, which occurs in cachexia [8]. Surprisingly, serum adiponectin levels were significantly lower in mice with CACS, as compared to mice without CACS (NCACS) and non-tumor bearing littermate controls (WT) (Figure 1C). These data suggest that CACS is associated with WAT dysfunction, including reduced production of adipokines like leptin and adiponectin.

**Figure 1.**
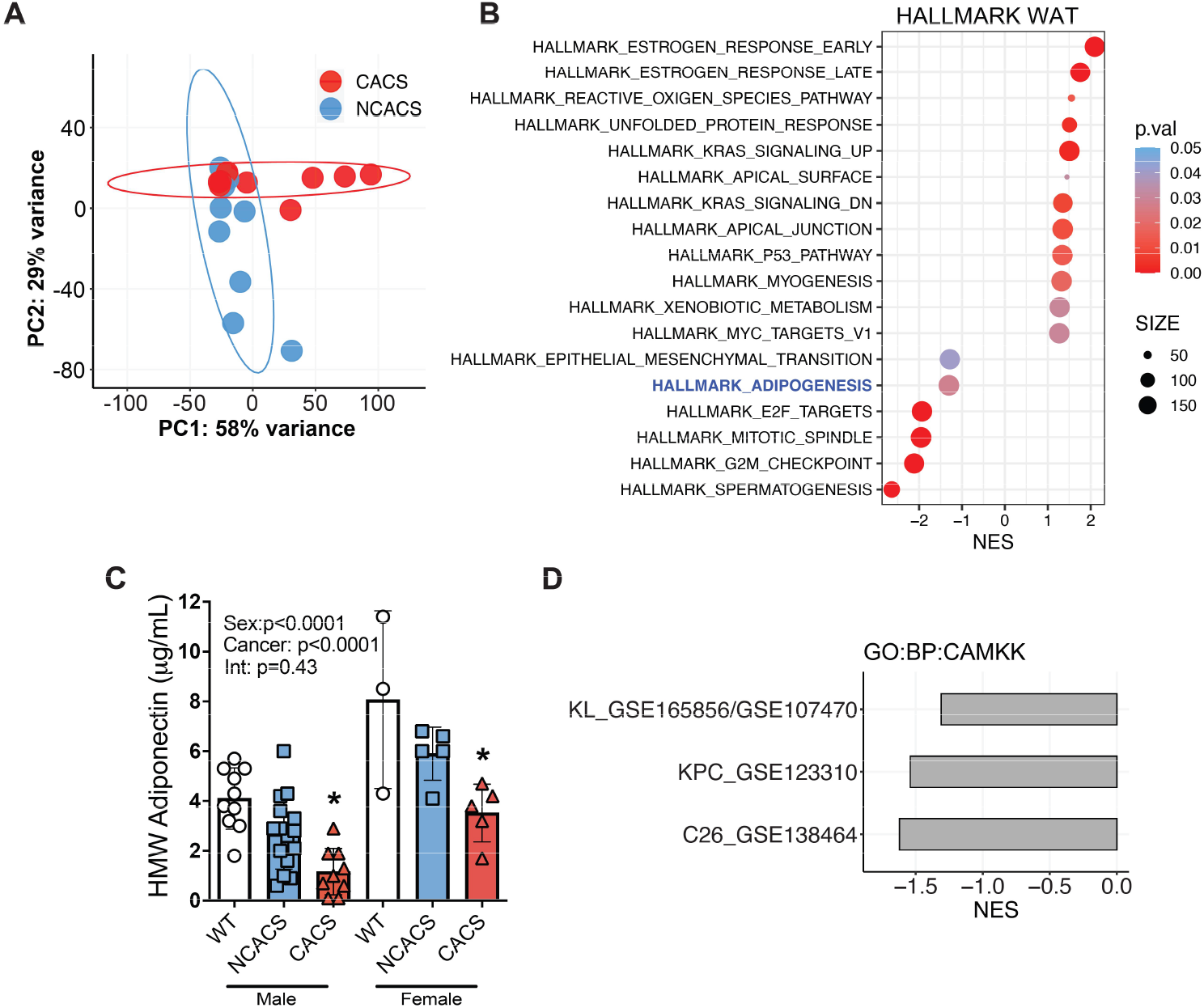
Cancer cachexia is associated with a decrease in serum adiponectin and a concomitant decline in AMPK/CaMKII signaling in skeletal muscle. A. Principal component analysis (PCA) of white adipose tissue (WAT) RNA-Seq samples showing distinct clustering between CACS (*n*=10) and NCACS (*n*=9) in the KL model. B. Gene Set Enrichment Analysis (GSEA) of samples in “A” showing the top 18 pathways differentially expressed in CACS using the Hallmark dataset. C. Serum levels of Adiponectin in wild type (WT, male=10, female=3) and tumor-bearing mice classified either as non-cachectic (NCACS, male=16, female=5) or cachectic (CACS, male=9, female=5). D. GSEA comparing CAMKK signaling pathway (GO:0061762) in the muscles of 3 different models of cancer cachexia. In GSE165856 and GSE107470, the gastrocnemius of CACS (*n*=10) KL mice are compared with those of NCACS (*n*=10). In, GSE123310 quadriceps of CACS (*n*=4) mice with Pancreatic ductal adenocarcinoma are compared to controls (*n*=3), and in GSE138464 the gastrocnemius of mice implanted with the C26 colon cancer (*n*=4) cell line are compared to controls (*n*= 6). Data information: Error bars stand for SD. p < 0.05, NES = Normalized Enrichment Score. The p-value in panel C was calculated using a two-way ANOVA followed by Sidak’s multiple comparison test.

Adiponectin is known to improve insulin sensitivity and stimulate glucose and fatty acid uptake into peripheral tissues, such as skeletal muscle, by activating AMP-activated protein kinase (AMPK) [9]. This activation of AMPK in muscle is at least partially dependent on binding of adiponectin to surface adiponectin receptors (AdipoR1/2), which signals a rise in intracellular calcium and auto-phosphorylation of Calcium/calmodulin-dependent protein kinase II (CaMKII) [10]. Using GSEA, we found transcriptomic evidence of reduced CaMKII activation in skeletal muscles from KL mice with CACS and two other commonly used animal models of cachexia (C26 colorectal cancer and KPC pancreatic cancer) (Figure 1D) [11, 12]. These results suggest that loss of adiponectin signaling occurs across various animal models of CACS.

### Rosiglitazone successfully restores circulating adiponectin in CACS mice with lung cancer

We hypothesized that restoring circulating adiponectin and re-activation of the CaMKII/AMPK pathway in skeletal muscle could be a viable strategy to preserve tissue mass and improve CACS in mice with lung cancer. The thiazolidinedione, rosiglitazone, is a PPAR-γ activator known to increase adiponectin synthesis and release from adipose tissue [13]. We induced lung tumors in KL mice as described previously [4, 6] and randomized them to receive a normal chow diet or a normal chow diet infused with rosiglitazone (100 mg/kg). Mice were serially monitored for changes in body weight and then we harvested key metabolic tissues and serum when they reached humane endpoint criteria. We found that rosiglitazone successfully restored levels of total and high molecular weight (HMW) adiponectin in the serum (Figure 2A-C). Adiponectin is composed of globular subunits and the HMW form is thought to be the biologically active form [14]. HMW adiponectin was approximately four times higher in the rosiglitazone group compared to the control (Figure 2B), and the ratio between HMW and total adiponectin was also significantly increased by rosiglitazone (Figure 2C). Since rosiglitazone is FDA-approved for use in patients with type 2 diabetes and known to modulate insulin sensitivity, glucose homeostasis, and lipid metabolism [9, 14], we measured circulating insulin, glucose, and triglyceride levels in the serum of our KL mice. No differences were found for insulin or glucose (Figure 2D-E). In keeping with the hypothesis that the drug would increase fatty acid oxidation in peripheral tissues, we found a decrease in circulating triglyceride levels with rosiglitazone (Figure 2F). Taken together, these data suggest that rosiglitazone can restore circulating adiponectin levels in mice with lung cancer without modulating whole-body glucose homeostasis.

**Figure 2.**
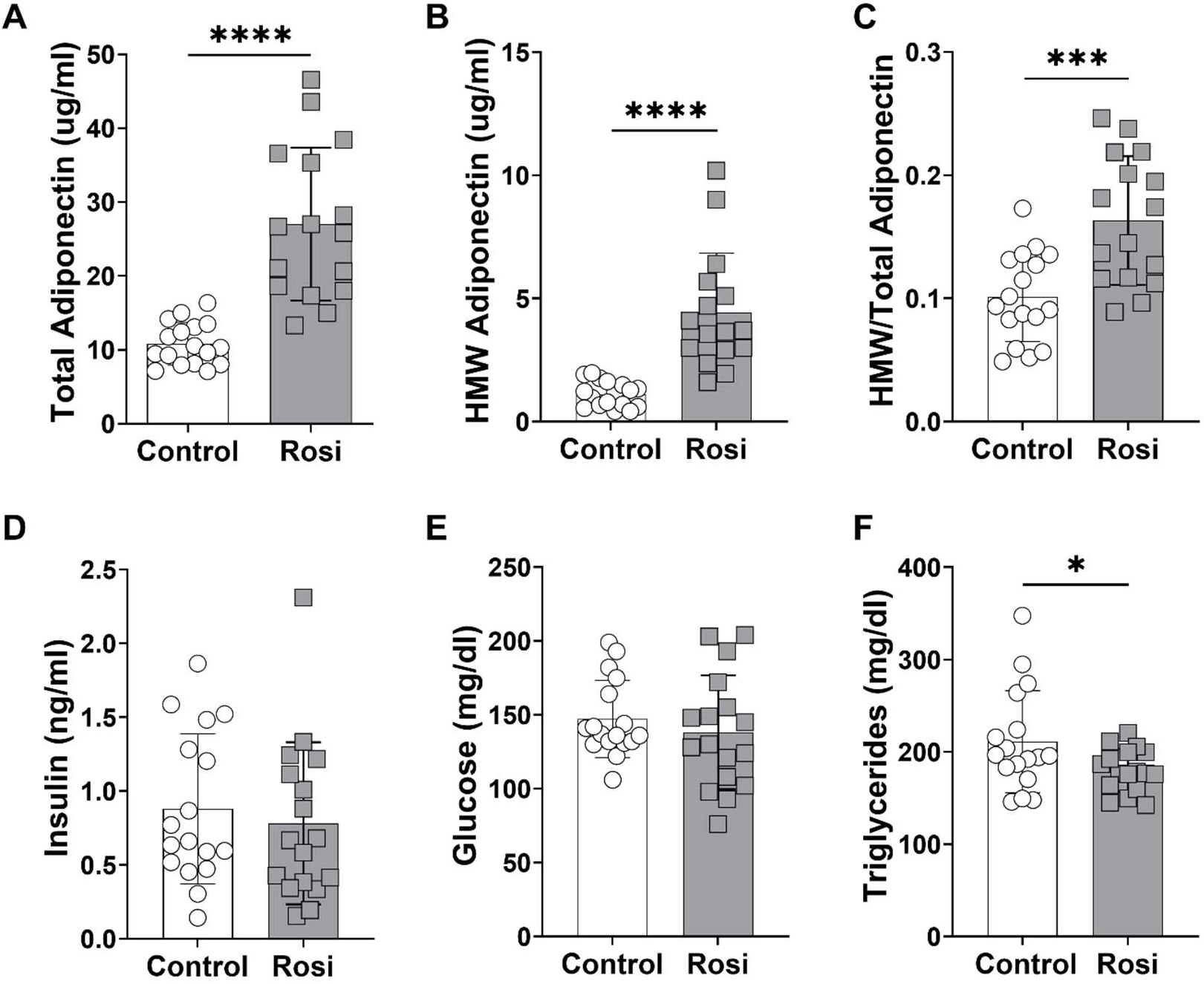
Serum analysis of cachectic mice treated with or without rosiglitazone. A-C. Rosiglitazone administration (Rosi) through *ad libitum* food intake (incorporated into regular chow, 100mg/kg) successfully restores total adiponectin, the physiologically active heavy molecular weight (HMW) adiponectin and the ratio of HMW to total adiponectin in cachectic mice (*n*=16-17 per group). D. No change in circulating insulin levels between Control and Rosi mice (*n*=17 per group). E. No change in blood glucose levels between Control and Rosi mice (*n*=17 per group). F. Rosiglitazone administration significantly reduced serum triglyceride levels in cachectic mice (*n*=17 per group). Data information: Error bars stand for SD. The p-value was calculated by an unpaired t-test. * denotes a p-value of <0.05, *** denotes a p-value of <0.001, and **** denotes a p-value of <0.0001.

### Rosiglitazone preserves skeletal muscle and adipose tissue mass

To investigate whether rosiglitazone improved features of CACS, we examined changes in body weight and tissue mass in the KL mice. There was a significant time-treatment interaction regarding the weight trajectory, indicating that rosiglitazone delayed weight loss (Figure 3A). These effects of rosiglitazone were independent of food intake, as both groups steadily decreased their intake over time without a significant impact of the treatment (Figure 3B). The preservation of body weight with rosiglitazone was associated with positive effects on peripheral tissues. Skeletal muscle mass (gastrocnemius) was significantly higher in the treatment group, as compared to the control group (Figure 3C). Furthermore, the rosiglitazone-treated mice had 97 % more WAT mass (Figure 3D) and 39 % more brown adipose tissue (BAT) mass (Figure 3E) than the controls. Lung mass is an excellent proxy for tumor burden in this model [4, 6, 15], and we found that both control and rosiglitazone-treated animals had similar amounts of tumor burden (Figure 3F). There was no difference in liver and spleen mass (data not shown). To summarize, the restoration of circulating adiponectin through rosiglitazone associates with delayed body weight loss and preserved muscle and adipose tissue mass, without affecting food intake or tumor growth.

**Figure 3.**
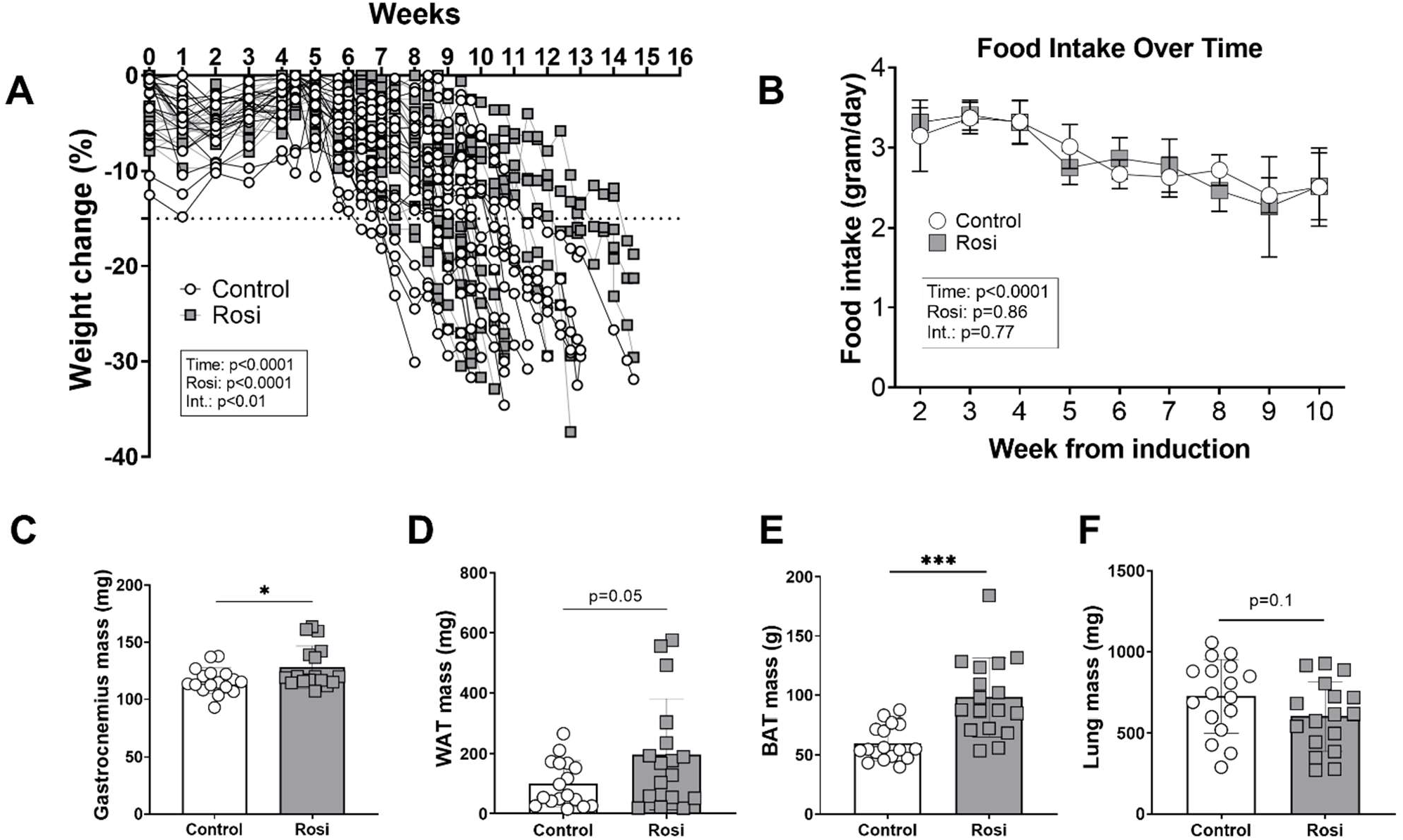
Restoring adiponectin through rosiglitazone results in preserved skeletal muscle and adipose tissue mass of cachectic mice. A. Both groups significantly reduced body weight over the 15-week intervention period (p<0.0001). There was a significant effect for the rosiglitazone intervention (p<0.0001) and a significant interaction effect between time and the rosiglitazone intervention (p<0.05) to affect body weight (*n*=17 per group). B. Food intake significantly declined over time (p<0.0001) but there was no effect of rosiglitazone (p=0.86) or an interaction between rosiglitazone and time (p=0.77). Food intake was calculated by dividing the total intake per cage by the number of animals per cage (*n*=4 cages per group). C-E. Skeletal muscle, white adipose tissue (WAT), and brown adipose tissue (BAT) were significantly preserved in the Rosi compared to the Control condition (*n*=17 per group). F. Rosiglitazone administration did not significantly affect lung weight in cachectic mice (*n*=17 per group). Data information: Error bars stand for SD. The p-value was calculated by a two-way ANOVA (body weight, food intake), and an unpaired t-test (tissue weights).

### Rosiglitazone activates AMPK activity and maintains anabolic signaling in skeletal muscle of mice with lung cancer

To learn how rosiglitazone preserved muscle mass at the molecular level, we performed immunoblot analysis on lysates from the gastrocnemius muscles. Of note, we decided to exclude the small subset of rosiglitazone-treated animals that maintained body weight exceptionally well throughout the entire experiment (Figure 3A) to avoid bias in our analysis. Previous reports have shown that adiponectin primarily exerts its effects on muscle through binding to adiponectin receptor 1 (AdipoR1) which in turn enables Ca^2+^ influx into the cytosol, leading to auto-phosphorylation of CaMKII at T286 [10]. In keeping with this logic, we found phospho-CaMKII (T286) to be upregulated 2.3-fold in rosiglitazone-treated animals, as compared to controls (Figure 4A). Similarly, increased AMPK activity is a hallmark of adiponectin signaling in skeletal muscle [9] and we found a 10-fold elevation in the levels of phosphorylated AMPK (T172) in the gastrocnemius of rosiglitazone treated animals (Figure 4B). The p38 MAPK is a known downstream target of adiponectin and AMPK in muscle [16, 17]. We measured phosphorylation of p38 at T180/Y182 and found a 50 % increase in the rosiglitazone group (Figure 4C). These data highly suggest that the restored adiponectin from the circulation is actively affecting signaling in the skeletal muscle.

**Figure 4.**
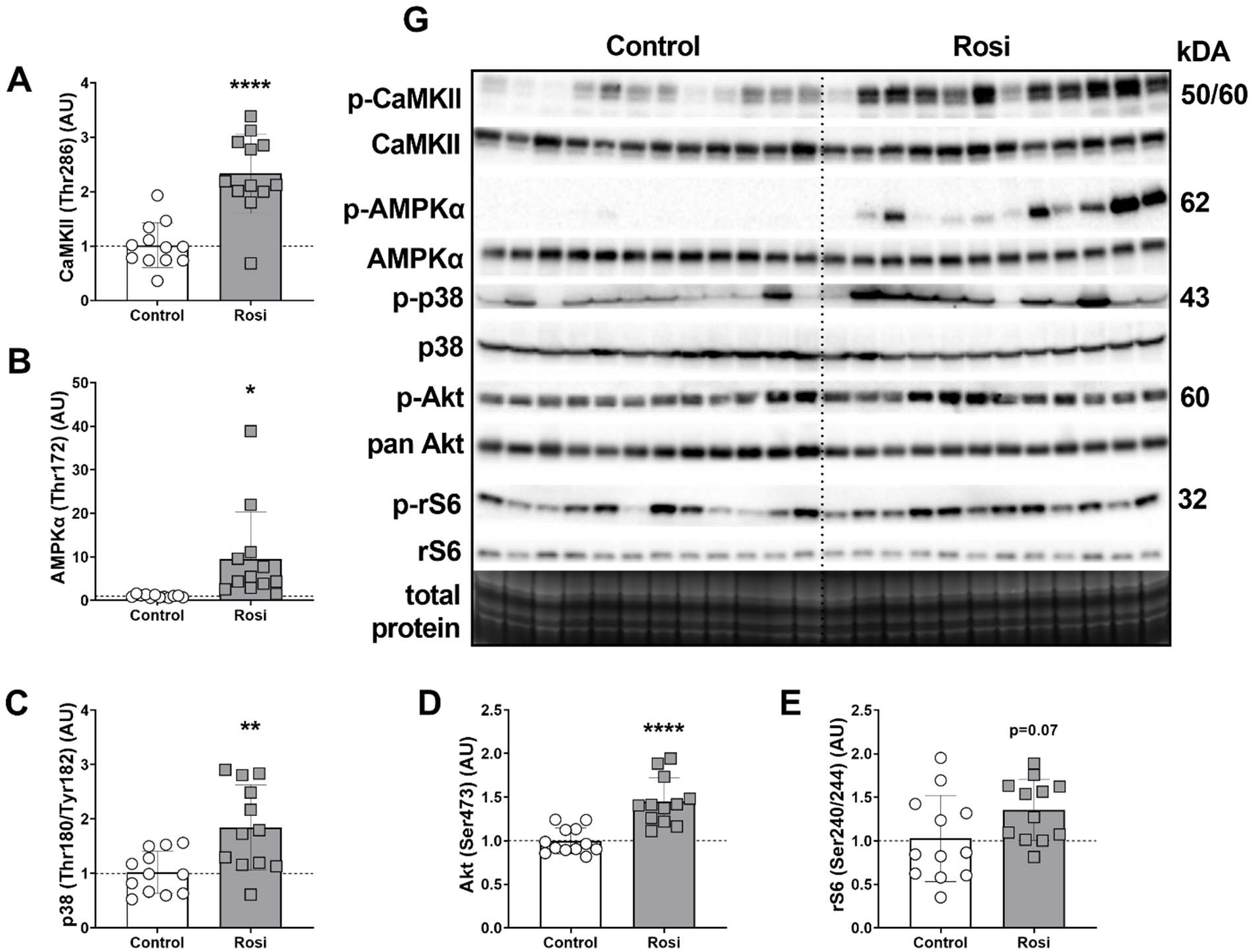
Restoring circulating adiponectin results in increased AMPK activity and improved maintenance of Akt signaling in skeletal muscle of cachectic mice. A-C. Western blot analysis of CAMKII, AMPK, and p38 signaling in gastrocnemius muscle of cachectic mice (*n*=12 per group). The bar graphs represent the ratio of phosphorylated to total protein levels. D-E. Western blot analysis of Akt activity and downstream signaling in gastrocnemius muscle of cachectic mice (*n*=12 per group). The bar graphs represent the ratio of phosphorylated to total protein levels. G. Whole membrane strips corresponding to the quantified western blot data from “A-F”. Data information: Error bars stand for SD. The p-value was calculated by an unpaired t-test. * denotes a p-value of <0.05, ** denotes a p-value of <0.01, and **** denotes a p-value of <0.0001.

Since muscle mass was improved with rosiglitazone treatment, we next investigated whether our intervention had positive effects on anabolic signaling. We found a 45 % increase in phosphorylation of Akt at S473 (Figure 4D), a site known to be targeted by mTORC2 following stimulation by mitogens like insulin [18]. Similarly, mTOR downstream target ribosomal protein S6 (rS6) showed an increasing trend with rosiglitazone (Figure 4E). Taken together, these data suggest that restoring adiponectin through rosiglitazone results in increased activity of CaMKII/AMPK/p38, but also a mildly positive effect on anabolic signaling through Akt/mTOR/S6. The latter offers a potential explanation for the improved maintenance of muscle mass during CACS with rosiglitazone since we previously found this pathway to be downregulated in cachectic KL mice [4].

### Adiponectin-receptor signaling activates AMPK and acutely improves anabolic signaling and protein synthesis in C2C12 myotubes

To further explore the relationship between adiponectin and anabolic signaling, we performed follow-up experiments in muscle cell culture. C2C12 cells were cultured with known activators of AMPK in the presence or absence of IL-6, which was added to simulate the inflammatory milieu present during CACS [11, 19]. First, we conducted a dose-response experiment to explore the acute effect of adiponectin on fully differentiated muscle cells. To mimic adiponectin, we used AdipoRon, a selective small molecule agonist of AdipoR1/2 *in vitro* and *in vivo* [20]. Three hours after adding AdipoRon to C2C12 myotubes, we found a 90 % increase in phospho-AMPK at T172 with the highest dose (Figure 5B). Interestingly, we found that with the same dose phospho-S6K1 at T389 was also significantly increased (Figure 5C). S6K1 is a known regulator of protein synthesis and required for muscle size maintenance and growth [21, 22]. To assess whether the changes in S6K1 would result in physiological effects on muscle protein synthesis in our experiment, we used the SUnSET method and added puromycin prior to collection. Puromycin is a t-RNA analog and gets incorporated into any newly synthesized peptide [23, 24]. By analyzing puromycin protein levels through western blot, one can obtain global protein synthesis rates for the time frame between application of puromycin to cells or tissue, and the point of collection [25, 26]. We found a significant increase in protein synthesis of ∼70 % for the moderate and high doses of AdipoRon. This result indicates that acute exposure of muscle cells to adiponectin receptor agonists results in a robust increase of AMPK activity and a concomitant increase in anabolic signaling as well as protein synthesis.

**Figure 5.**
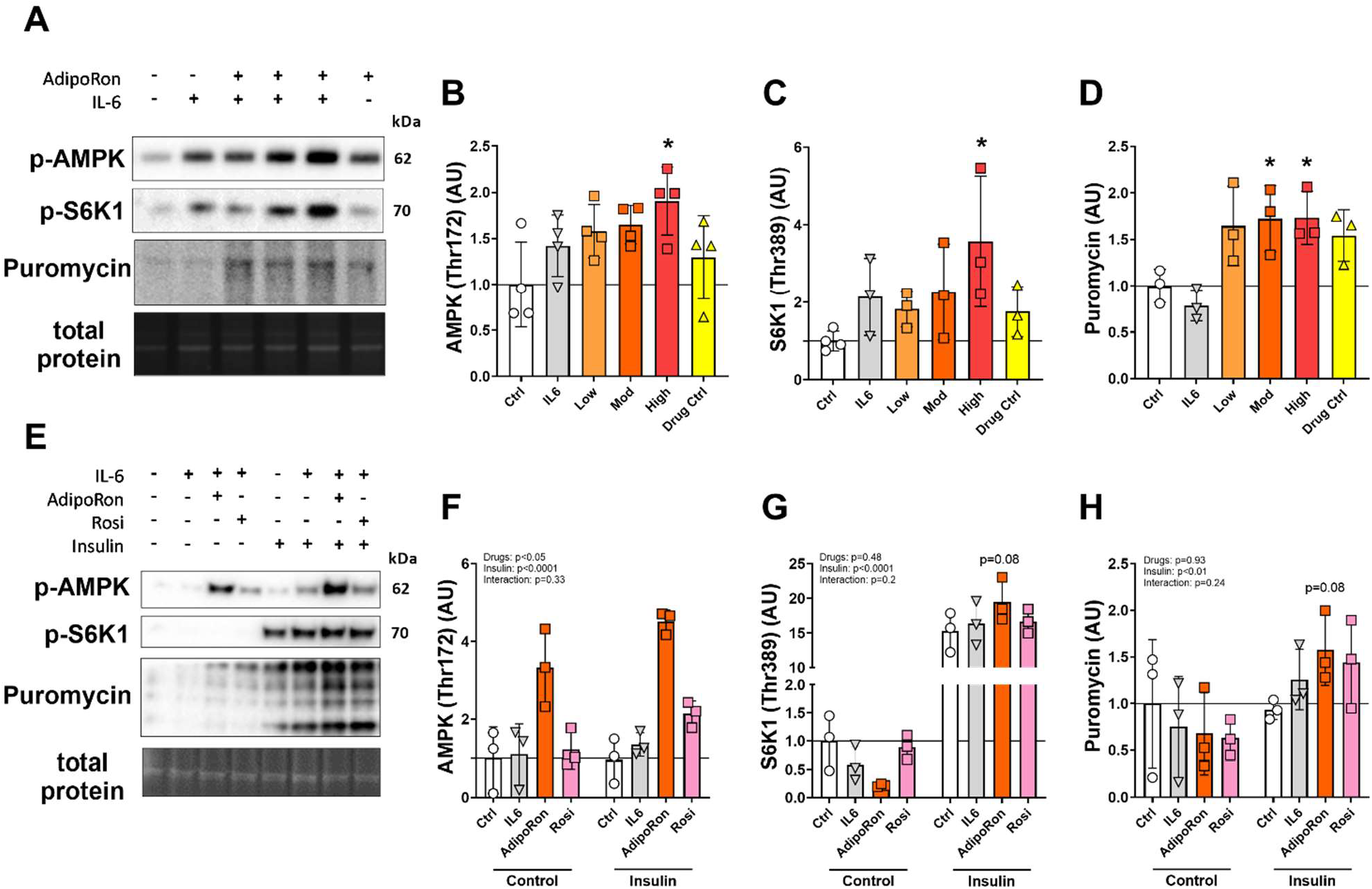
Adiponectin activates AMPK and acutely improves muscle protein synthesis in the presence of IL-6, while chronic effects appear insulin-dependent. A-D. Western blot analysis of AMPK activity, anabolic signaling, and protein synthesis in C2C12 cells after an acute dose-response experiment (*n*=4 per group). Cells were treated with increasing doses of adiponectin (10, 25, 50µM) together with IL-6 (10ng/mL) for 3 hours. The control condition was untreated, the IL-6 condition received only IL-6 (10ng/mL), and the Drug Ctrl group received only adiponectin (25µM). “A” shows representative pictures of the graphs from “B-D”. Phosphorylated levels of AMPK (B), S6K1 (C), and puromycin (D) were normalized to total protein content per lane. E-H. Western blot analysis of AMPK activity, anabolic signaling, and protein synthesis in C2C12 cells after chronic exposure to adiponectin or rosiglitazone (*n*=3 per group). Cells were treated with adiponectin (25µM) or rosiglitazone (10µM) together with IL-6 (10ng/mL) for 24 hours. Thirty minutes before collection, half of the cells were treated with 100nM insulin. The Ctrl group was untreated, the IL- 6 group only received IL-6 (10ng/mL). “E” shows representative pictures of the graphs from “F-H”. Phosphorylated levels of AMPK (F), S6K1 (G), and puromycin (H) were normalized to total protein content per lane. Data information: Error bars stand for SD. The p-value was calculated by a one-way ANOVA and Dunnett’s multiple comparison test (acute experiment, Figure 5A-D), and a two-way ANOVA and Sidak’s multiple comparison test (chronic experiment, Figure 5E-H). * denotes a p-value of <0.05.

To understand whether these results could be replicated in a chronic scenario, and to learn whether the effect of adiponectin was insulin-dependent, we performed additional experiments in C2C12 myotubes lasting 24 hours. Cells were treated with either IL-6, IL-6, and the moderate dose of AdipoRon, or IL-6 and rosiglitazone. Thirty minutes prior to the 24-hour mark, half of the cells were treated with insulin. In the absence of insulin, we found that AdipoRon, but not rosiglitazone, increased phospho-AMPK at T172, resulting in a significant main-effect for differences between the drugs (Figure 5F). In line with the known interactions between adiponectin and insulin signaling [9, 27], we found that insulin exacerbated the effect of AdipoRon and rosiglitazone on AMPK (Figure 5F). Insulin also robustly activated phospho-S6K1 (T389), with no significant differences when AdipoRon or rosiglitazone were added (Figure 5H). Of note, there was a trend for AdipoRon to increase the abundance of phospho-S6K1 (p=0.08). Lastly, we investigated the effect of the signaling changes on protein synthesis in the cells. In keeping with the phospho-S6K1 data, we found a main-effect for insulin to increase protein synthesis, with the only trend for a significant increase between baseline and the insulin-addition being in the AdipoRon group (p=0.08) (Figure 5I). Our results suggest that stimulation through the adiponectin receptor is likely the predominant pathway through which AMPK activation and muscle anabolism were maintained in the rosiglitazone-treated mice and that the chronic effects of adiponectin on protein synthesis were at least partially insulin dependent.

## Discussion

CACS is a catabolic wasting syndrome associated with the loss of skeletal muscle and adipose tissue mass. In this study, we find evidence of transcriptional and physiologic dysfunction in adipose tissue that directly affect skeletal muscle. Typically, there is an inverse relationship between adipocyte size and adiponectin levels. When adipocyte size increases in the setting of obesity, there is a decrease in adiponectin levels, which can contribute to the development of insulin resistance and other metabolic disorders [28]. When adipocytes decrease in size in the setting of calorie restriction, adiponectin levels rise and support fatty acid oxidation and insulin sensitivity in other organs. The fact that the paradoxical combination of adipocyte atrophy and low adiponectin levels during CACS suggests adipocyte dysfunction in line with the phenotype observed in humans with lipodystrophy, another condition associated with severe metabolic dysfunction [29].

The rise in circulating adiponectin level is likely a key mediator of the rosiglitazone treatment. While we cannot rule out adiponectin-independent effects, the increase in the abundance of adiponectin signaling components such as CAMKII, AMPK, and p38 in the skeletal muscle suggests adiponectin is more active in the rosiglitazone-treated muscle. Also, the cell culture data support a direct role of adiponectin on skeletal muscle protein synthesis. We find that stimulation of the adiponectin receptors with AdipoRon can acutely activate anabolic signaling and protein synthesis in a dose-dependent manner. These effects were not accomplished by rosiglitazone alone suggesting this agent does not act directly on skeletal muscle. In support of our data, a recent study has shown that recombinant adiponectin activates Akt and S6K1 signaling in C26-conditioned myotubes and that this treatment resulted in improved cell size [30].

Our data suggest a mechanistic link between adiponectin/CaMKII/AMPK and anabolic signaling through Akt/mTOR/S6K1 in skeletal muscle. A recent study has shown that AMPK may directly phosphorylate and activate mTORC2, which helps promote cell survival under acute energetic stress [31]. Indeed, in our study, we found a modest increase in phospho-Akt at serine S473, which is thought to be primarily phosphorylated by mTORC2. This stimulation may mediate the modest, positive effects on muscle anabolism and body weight, especially since we previously found that Akt/mTOR/S6 signaling is impaired in cachectic compared to non-cachectic KL mice [4]. In keeping with this idea of a positive relationship between AMPK and anabolic signaling, a recent study found that the AMP-analog AICAR successfully rescued anabolic signaling in C2C12 cells treated with inflammatory cytokines, and maintained muscle mass in mice with cachexia [32]. Comprehensive work in wild-type and muscle-specific AMPK-null mice with lung cancer indicates that this increase in tissue anabolism is at least partially mediated by the insulin-sensitizing effects of AMPK [33, 34]. A study that focused on the role of AMPK in adipose tissue found that an AMPK-stabilizing peptide was able to ameliorate tissue wasting in colorectal-and lung cancer [35]. Thus, it seems plausible that activating AMPK may have properties that enable the maintenance of anabolic signals, protein synthesis, and tissue mass, particularly in catabolic scenarios.

This study is the first to show that thiazolidinediones can improve CACS in mouse models of lung cancer. Our results agree with others who have used these drugs to preserve body weight in animal models with CACS [36-40]. Asp et al. found that rosiglitazone could prevent weight loss and preserve WAT mass in mice bearing C26 allografts [37, 38]. The beneficial effects were found to be due to a delayed onset of anorexia, increased insulin sensitivity, and a suppression of proteolysis-promoting genes. Rosiglitazone also reduced weight loss in rats injected with Yoshida AH-130 hepatoma tumor cells [39]. In this setting, the beneficial effects on body weight were also primarily due to the preservation of adipose tissues as muscle mass was not protected. Interestingly, they observed a reduction in the activity of the proteasome system in skeletal muscle despite no difference in tissue mass. Another thiazolidinedione, pioglitazone, was assessed in Wistar rats inoculated with Walker 256 tumor cells [36]. Here, the drug delayed weight loss and anorexia, protected adipose tissue mass, and improved overall survival. However, the treatment also slowed tumor growth, which makes it difficult to determine if the beneficial effects on the host were due to less tumor growth or the drug. Together, these results suggest that thiazolidinediones could be an effective therapy to preserve body weight across tumor types, which may enable a prolonged duration of anti-cancer therapy.

Our study has a few limitations. We did not assess physical activity or muscle performance to confirm that the maintenance of gastrocnemius mass in our experiment was associated with functional improvements in strength or endurance capacity. In the setting of sarcopenia, strength is lost at a faster rate than muscle mass, and physical performance appears to be a relevant predictor of health [41, 42]. However, the amount of muscle mass independently predicts the tolerability of anti-cancer therapy, quality of life, and life expectancy in cachexia patients [43-45]. Therefore, our assessments of muscle and adipose tissue mass explain the attenuation of body weight loss in our model while also addressing a clinically relevant endpoint in human cachexia. Another limitation is that we did not directly measure insulin sensitivity, which may contribute to the anabolic signaling in skeletal muscle. Thiazolidinediones are known to improve insulin sensitivity, particularly in the context of diabetes, and other mouse models of cachexia have shown that rosiglitazone increases insulin sensitivity [38]. In keeping with these data, our 24-hour cell culture experiment suggested that insulin is at least partially involved in mediating the effects of rosiglitazone and adiponectin in our model. Therefore, we suspect that an improvement in insulin sensitivity is contributing to the overall benefits of rosiglitazone in our study. Despite these limitations, our data reveal the previously unappreciated effect of rosiglitazone and adiponectin on anabolic processes and tissue maintenance in mouse models of lung cancer. These findings support future clinical trials using thiazolidinediones in patients with lung cancer at high risk for cachexia.

## Methods

### Cachexia model and lung tumor induction

*Kras^G12D/+^;Lkb1^f/f^* (KL) mice have been previously described [46]. Mice were housed in a 12-h light/dark cycle and 22°C ambient temperature and had free access to normal chow (PicoLab Rodent 20 5053; Lab Diet) or rosiglitazone-containing chow (Modified Lab Diet 5053 with 100 ppm Rosiglitazone, cat no. 5WXY, Adooq Bioscience, cat no. A10807) as well as drinking water. Tumors were induced in adult (9- to 18-wk-old) male mice via intranasal administration of 75 μL of Optimem containing 1 mM CaCl2 and 2.5 × 10^7^ pfu of Adenovirus CMV-Cre (Ad5CMV-Cre) purchased from the University of Iowa Gene Transfer Vector Core (Iowa City, IA). A detailed description of the cachectic phenotype in these mice, its time-course, and its severity have been published previously [4, 6]. Mice were euthanized if they reach the following humane endpoints: ≥30% loss from peak body weight or poor body composition score (<2).

### Study approval

All animal studies were approved and maintained as approved by the Institutional Animal Care and Use Committee (IACUC) of Weill Cornell Medicine under protocol number 2013-0116.

### Tissue Collection Protocol

Prior to euthanasia, glucose was measured from tail vein blood using a handheld point-of-care glucose meter (OneTouch). Euthanasia was performed using CO2 asphyxiation. Following euthanasia, whole blood was collected via cardiac puncture and placed into serum separator tubes on ice. Subsequently, the whole liver was removed, weighed, and frozen in liquid nitrogen. The white and brown adipose tissues, lungs, spleen, and gastrocnemius muscles were dissected, weighed, and flash-frozen in liquid nitrogen or fixated in paraformaldehyde. All tissues were subsequently stored at −80°C until further processing.

### RNA sequencing analysis

RNASeq analysis was performed in previously published datasets GSE165856, GSE107470, GSE123310, GSE138646. Differential expression analysis, batch correction, and principal component analysis (PCA) were performed using R Version 4.2.2 and DESeq2 (v.1.38.3). Gene set enrichment analysis (GSEA) analysis was performed using GSEA (v4.3.2) JAVA based application. Gene sets for GSEA were obtained from the Mouse MSigDB Collections.

### Serum hormones and metabolites

Blood was collected in SST tubes, centrifuged at 10,000 × g for 10 min at 4°C, and serum was stored at −20°C. Serum adiponectin (ALPCO Diagnostics, cat no. 47-ADPMS-E01), insulin (Crystal Chem cat. no 90080), and triglycerides (LiquiColor, cat no. 2100-225) were determined using commercially available kits, per the manufacturer’s instructions.

### Cell culture experiments

C2C12 myoblasts were cultured in growth media (high glucose Dulbecco’s modified Eagle medium [DMEM, Life Technologies cat no. 11965118], 10 % fetal bovine serum, 0.1 % penicillin[Gibco, 15140122]) until they reached 95–100 % confluence at which time, they were shifted into differentiation media (DM; high glucose DMEM, 2 % horse serum, 0.1 % penicillin) to promote the differentiation and fusion of cells to form myotubes. All experiments were conducted on fully formed myotubes 5 days after the introduction of DM. For the dose-response experiment using AdipoRon (ChemCruz, cat no. sc-396658A), a low (10 µM), moderate (25 µM), and high (50 µM) dose were applied to the cells 3 h prior to collection. All three groups were co-treated with IL-6 (10 ng/mL, BioLegend cat no. 575706). We included one control group that was treated with DMSO, one control group with only IL-6, and one with only the moderate dose of AdipoRon. For the chronic experiments, we incubated C2C12 myotubes for 24 h with DMSO, IL-6 (10 ng/mL), AdipoRon (25 µM), and IL-6, or rosiglitazone (10 µM) and IL-6. 30 minutes prior to collection of the cells, insulin (100 nM, Sigma Aldrich # I9278-5ML) was added to half of the cells to explore insulin-dependency of the signaling events. For all cell culture experiments, puromycin (45 μM, Research Products International cat no. P33020-0.1), was added to the wells 5 min before collection to measure muscle protein synthesis via the SUnSET method [47]. For western blot analysis, 24 or 48 well plates were washed with ice-cold PBS three times before 75 μL of 1×LSB was directly added to each well and the samples were transferred to a fresh tube. Samples were then briefly sonicated before boiling at 100°C and stored at −30°C until further analysis.

### Western blotting

Immunoblotting was carried out as described previously [48]. Briefly, for the mouse gastrocnemius samples 10 to 40 μg of protein per lane were loaded. For the cell culture experiments, 10 to 15 μL of sample were loaded per lane. All samples were run for 40 min at 200 V before transfer onto polyvinylidene fluoride or nitrocellulose membrane in an ice-cold transfer buffer at 100 V for 30 min. Membranes were blocked in 1% fish skin gelatin dissolved in Tris-buffered saline with 0.1% Tween-20 for 30 min, or air-dried, and probed with the primary antibody overnight. The following antibodies were used: Cell Signaling (Cell Signaling Technology, Danvers, MA): CaMKII (S286) (No. 12716), pan CaMKII (No. 4436), AMPK (T172) (No. 2535), AMPK (No. 2532), p38 (T180/Y182) (No. 4511), p38 (No. 9212), Akt (S473) (No. 4060), pan Akt (No. 4691), ribosomal protein S6 (Ser240/244) (No. 5364), S6K1 (Thr389) (No. 9205; lot 16); Millipore Sigma (Merck Group): puromycin (No. MABE343). Protein levels were normalized to the unphosphorylated version of the protein or the total protein content per lane as assessed via stain-free TGX gel technology (Bio-Rad, Hercules, CA, USA) [49]. Image acquisition and band quantification was performed using the ChemiDoc MP System and Image Lab 5.0 software (Bio-Rad).

### Statistics

Statistical analysis was carried out using Prism 9.5.1 (GraphPad Software). Individual data points are provided for every graph and assay with exception of the RNAseq data. The mean ± SD is displayed in addition to the individual data for all data pertaining to the rosiglitazone intervention and the cell culture data. The statistical test applied was dependent on the analysis performed and is indicated in each legend. An unpaired t-test was used to compare serum hormone and substrate levels (Figure 2A-F), tissue weights (Figure 3B-G), and protein levels in skeletal muscle (Figure 4A-E). A two-way ANOVA was performed to detect main effects for the rosiglitazone intervention, time, and any potential interactions between rosiglitazone and time (Figure 3A). A one-way ANOVA and Dunnett’s multiple comparison test were used to analyze the dose-response experiments in cell culture (Figure 5A-D). A two-way ANOVA was performed to determine main effects of the drug interventions, insulin, and any interactions between the drugs and insulin in our chronic cell culture experiment (Figure 5E-H). Sidak’s multiple comparison test was run post-hoc to investigate individual differences in the response to insulin between the groups. A p-value of <0.05 was deemed statistically significant, values between 0.05 and 0.1 are referred to as trends.

## Acknowledgements

This work was supported by NIH K08CA230318 (M.D.G.), the 2020 AACR-AstraZeneca Lung Cancer Research Fellowship Grant Number 20-40-12-DANT (E.D.), and was delivered as part of the CANCAN team supported by the Cancer Grand Challenges partnership funded by Cancer Research UK (CGCATF- 2021/100022) and the National Cancer Institute (1 OT2 CA278685-01).

## Author contributions

HTL designed and executed the cell culture experiments, performed all in vivo and vitro immunoblot analyses, contributed to the serum assays, performed data analyses, and wrote the manuscript. SR designed and performed the animal studies, performed tissue collections and serum analyses, and contributed to the cell culture experiments. ED performed data mining and RNAseq analyses. EW and TJ contributed to the drafting of the manuscript. MDG designed the animal studies, helped analyze the data and drafted the manuscript. All other authors were involved in contributing to the animal experiments and a priori data collection that helped guide the project.

